# More than a feeling: scalp EEG and eye correlates of conscious tactile perception

**DOI:** 10.1101/2021.10.31.466706

**Authors:** Mariana M. Gusso, Kate L. Christison-Lagay, David Zuckerman, Ganesh Chandrasekaran, Sharif I. Kronemer, Julia Z. Ding, Noah C. Freedman, Percy Nohama, Hal Blumenfeld

## Abstract

Understanding the neural basis of consciousness is a fundamental goal of neuroscience. Many of the studies tackling this question have focused on conscious perception, but these studies have been largely vision-centric, with very few involving tactile perception. Therefore, we developed a novel tactile threshold perception task, which we used in conjunction with high-density scalp electroencephalography and eye-metric recordings. Participants were delivered threshold-level vibrations to one of the four non-thumb fingers, and were asked to report their perception using a response box. With false discovery rate (FDR) mass univariate analysis procedures, we found significant event-related potentials (ERP) including bilateral N140 and P300 for perceived vibrations; significant bilateral P100 and P300 were found following vibrations that were not perceived. Significant differences between perceived and not perceived trials were found bilaterally in the N140 and P300. Additionally, we found that pupil diameter and blink rate increased and that microsaccade rate decreased following vibrations that were perceived relative to those that were not perceived. While many of the signals are consistent with similar ERP-findings across sensory modalities, our results indicating a significant P300 in not perceived trials raise more questions regarding P300’s perceptual meaning. Additionally, our findings support the use of eye metrics as a measure of physiological arousal as pertains to conscious perception, and may represent a novel path toward the creation of tactile no-report tasks in the future.

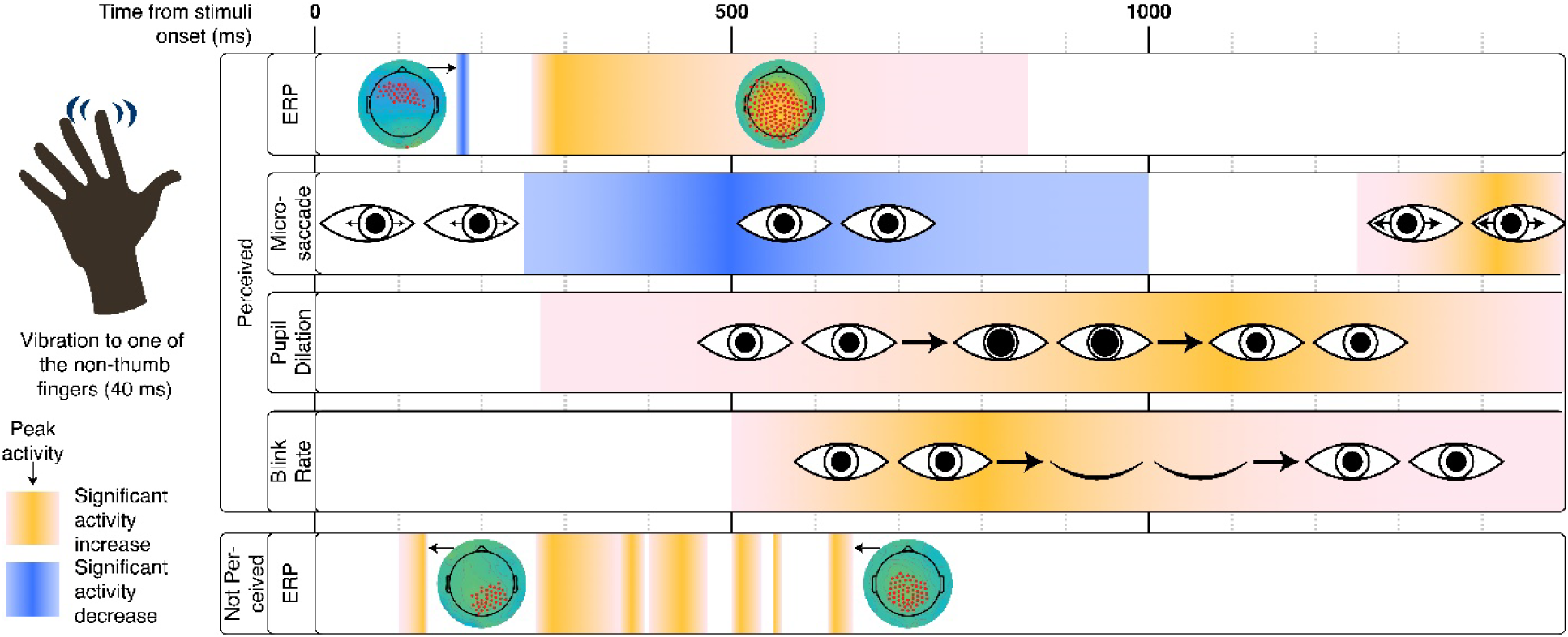

**Highlights:**

- A novel tactile perceptual threshold task yields robust behavioral results
- Event-related potentials differ according to perception status
- P300 is observed in both perceived and not perceived trials
- Blink rate, pupil diameter, and microsaccades differ across trial conditions

## 1 Introduction

One of the biggest challenges in modern science is to understand the neural basis of consciousness (Miller, 2005). Most studies have approached this through the lens of perceptual processing, using a combination of perceptual behavioral tasks coupled with various brain recording techniques, such as scalp and intracranial electroencephalography (EEG), magnetoencephalography (MEG), functional magnetic resonance imaging (fMRI) and direct neural recordings in both humans and animal models (Del Cul, Baillet, & Dehaene, 2007; Fukuda & Matsunaga, 1983; Li, Hill, & He, 2014; Pitts, Metzler, & Hillyard, 2014; Wyart & Tallon-Baudry, 2008). Although there are several conflicting theories about what gives rise to consciousness itself, studies spanning recording techniques and behavioral paradigms find characteristic activity in sensory areas followed by widespread activity in higher-level associative cortical regions, including frontal and parietal cortices. Studies on visual perception have dominated the field, and although auditory perceptual studies have made inroads, the number of studies examining other senses, including tactile (Eimer, Forster, & Van Velzen, 2003; Kida, Wasaka, Nakata, Akatsuka, & Kakigi, 2006; Schubert, Blankenburg, Lemm, Villringer, & Curio, 2006) and olfaction (Abbasi et al., 2020; Kim, Bae, Jin, & Moon, 2020) still lags far behind; though, notably, there is an extensive literature on pain perception that stands somewhat apart (Babiloni et al., 2001; Buchgreitz, Egsgaard, Jensen, Arendt-Nielsen, & Bendtsen, 2008; Douros, Karrer, & Rosenfeld, 1994; Egsgaard et al., 2012; McDowell et al., 2006; Truini et al., 2004). An expanded and rigorous study of the neural basis of consciousness across all sensory modalities is necessary to truly understand whether there are common mechanisms of conscious perception.

The existing literature on somatosensory perception has focused largely on masking or oddball paradigms (Eimer et al., 2003; Kida et al., 2006; Schubert et al., 2006); or studies involving multiple sensory modalities (Eimer et al., 2003; Montoya & Sitges, 2006). Although these studies provide valuable insight into perceptual processing, such studies almost always present *different* and/or *additional* masking stimuli to control whether or not a target is perceived. The potential confound of differences in the stimulus itself begs the question: was the observed brain activity across masked and unmasked conditions different because of the perceptual difference or because of differential stimuli (e.g., two stimuli in a masked condition vs. one in an unmasked one)? The use of a threshold detection task eliminates this potential confound, because identical (or functionally identical, as is the case with a perceptual threshold that changes over time) stimuli are presented: only the percept changes, either perceived or not perceived. Threshold detection tasks have successfully been used in vision (Herman et al., 2019; Kronemer et al., 2021; Pins & Ffytche, 2003; Ress & Heeger, 2003; Wyart & Tallon-Baudry, 2008) and audition (Christison-Lagay et al., 2018; Colder & Tanenbaum, 1999). To our knowledge, only one such tactile task has been published, but this task required participants to immediately move the stimulated finger (Palva, Linkenkaer-Hansen, Näätänen, & Palva, 2005); the immediate behavioral response complicates the interpretation of brain activity, as perceptual and motor components happen essentially at the same time.

Despite these caveats, a picture of the event-related potentials (ERPs) modulated by tactile perception has begun to emerge from previous studies. Forster, Tziraki, and Jones (2016), Kida et al. (2006) and Schubert et al. (2006) suggest that the P100 and N140 are the earliest signals modulated by attention, signals also pointed out by Schubert et al. (2006) as linked to conscious awareness.

While early potentials are often associated with specific sensory modalities, the P300 is commonly reported across modalities. However, despite its prevalence, its interpretation is controversial: it is debated whether the P300 is the result of processes necessary for awareness (Ye, Lyu, Sclodnick, & Sun, 2019) or attention (Pitts, Metzler, et al., 2014), or whether it is the result of post-perceptual processing (Cohen, Ortego, Kyroudis, & Pitts, 2020; Koivisto, Salminen-Vaparanta, Grassini, & Revonsuo, 2016; Kronemer et al., 2021; Muñoz, Reales, Sebastián, & Ballesteros, 2014; Railo, Koivisto, & Revonsuo, 2011).

Interpretation of electrophysiological signals of perception is further complicated by the use of behavioral report in nearly all perceptual tasks. Because the act of reporting recruits additional cognitive processes, there is an increasing call in the field to move toward tasks that require no perceptual report. When subjects are required to report whether or not they had perceived stimulus, this report recruits additional cognitive processes such as the retention of percepts in working memory, the preparation of a motor plan, etc. This poses a particular challenge for perceptual threshold tasks: although these tasks are very useful in identifying brain activity caused by perceptual differences (as opposed to changes correlated with physically different stimuli), the only difference between trials are, in fact, the participant’s perception, which must be read-out in some fashion. Therefore, development of covert measures of conscious perception are particularly important. One promising avenue of study has proposed using pupil diameter, blink, and microsaccade rates to covertly measure changes in physiological arousal, which in turn correlate with changes in cognitive engagement and perception (Eckstein, Guerra-Carrillo, Singley, & Bunge, 2017; Einhauser, Koch, & Carter, 2010; Kang & Wheatley, 2015; Kronemer et al., 2021; Laeng & Endestad, 2012; Piquado, Isaacowitz, & Wingfield, 2010). As with other methodologies, eye metrics have most frequently been used with visual paradigms; however, isolated studies have shown differences in pupil diameter (C. R. Lee & Margolis, 2016; van Hooijdonk et al., 2019), and microsaccade rate (Badde, Myers, Yuval-Greenberg, & Carrasco, 2020; Dalmaso, Castelli, Scatturin, & Galfano, 2017) associated with tactile perception.

Here, we present findings using a novel tactile threshold task, which was conducted with concurrent high-density scalp EEG and eye metric recordings. We find that perceived and not perceived trials, presented at an individual’s tactile perceptual threshold, exhibit differences in ERPs, pupil diameter, and blink and microsaccade rates. To our knowledge, this is the first time that a tactile threshold task has been performed using both pupillometry and high-density scalp EEG to help elucidate the underlying mechanisms of consciousness.

## 2 Materials and Methods

### 2.1 Participants

Twenty-six participants were recruited to participate in the task. From those, 10 participants completed the behavioral task with simultaneous high density electroencephalography (hdEEG) alone, and 16 with hdEEG concurrent with eye metrics. Of these, two participants were entirely excluded from analysis due to poor behavioral performance; four were excluded from eye metric analyses and one from hdEEG analysis due to inadequate data collection. Data analysis of hdEEG signals was completed for 23 participants (10 male; 6 left-handed); analysis of eye metrics was completed for 10 participants (4 male; 3 left-handed). All experimental procedures were approved by the Yale University Institutional Review Board and all participants provided written informed consent.

### 2.2 Task design

The behavioral task tested tactile conscious perception using a vibration delivered to the pad of one of the four non-thumb fingers (Fig. 1B). One hand was designated as the stimulus receiving hand and the other hand as the response hand; the hand selection was counterbalanced across individuals. During the course of artifact rejection and exclusion criteria (see below), more individuals receiving stimulation to the left hand ended up being excluded, so in total the data shown include 15 participants who received stimulation of the right hand and 9 participants who received stimulation of the left hand. Vibrating tactors (C-2 tactor, Engineering Acoustics, Inc.) were secured to each of a participant’s fingers using adjustable foam straps and a custom-made positioning template. Straps were color-coded to correspond to their counterpart button on a four-button response box (Current Designs, Inc) that was controlled by the hand contralateral to the hand receiving stimuli.

**Figure 1:**
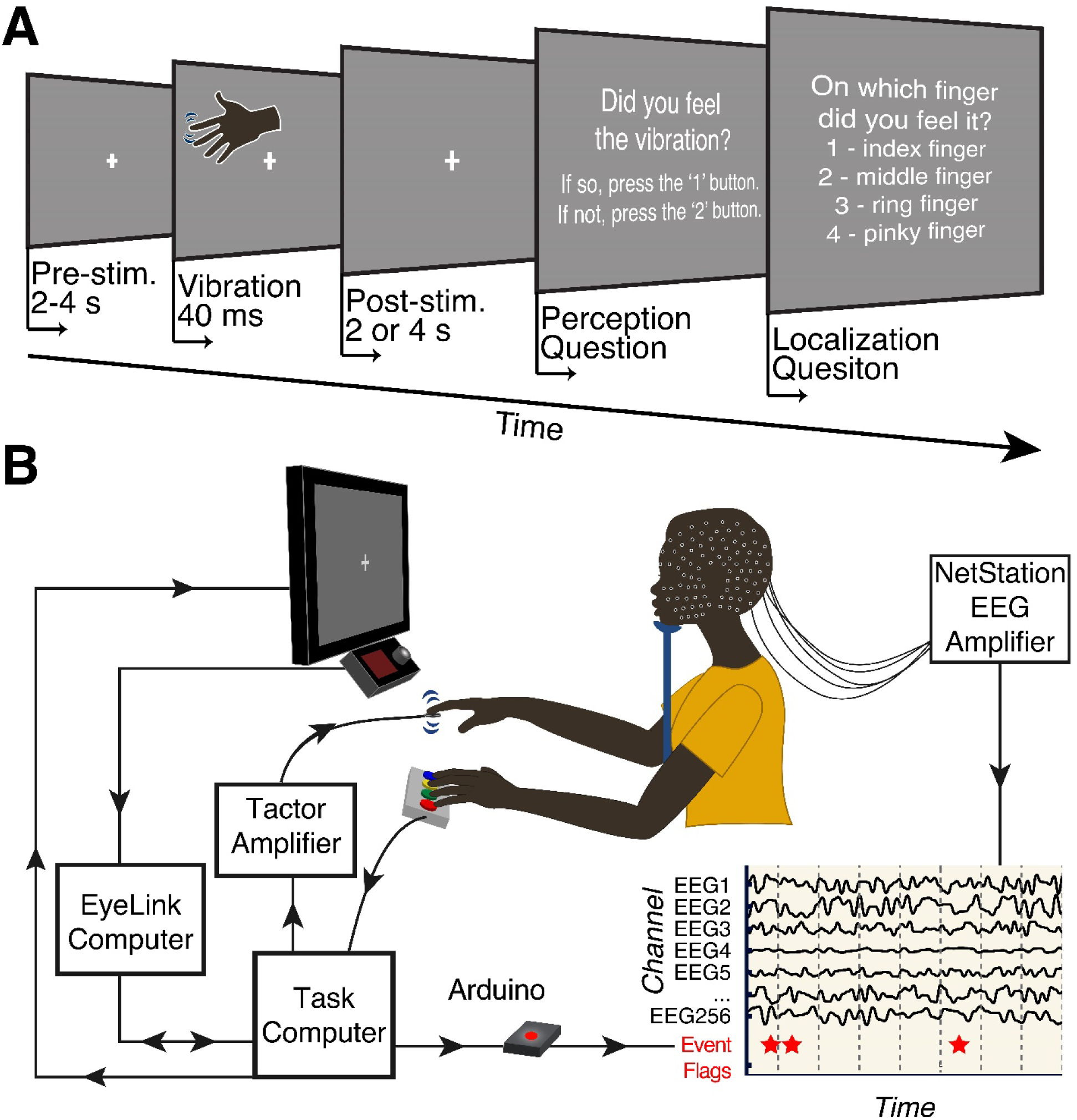
Tactile threshold task and experimental set-up. (**A**) Threshold tactile task for a single trial. Trials began with a randomly jittered pre-stimulus duration of 2-4 s of a gray screen with a white fixation cross, which was followed by a 40 ms 200 Hz sinewave vibration presented to one of a participant’s fingers (index, middle, ring or pinky) at the participant’s tactile threshold. After a post-stimulus delay of 2 or 4 s, participants were prompted (on-screen) to answer two forced choice questions regarding 1) whether they felt a stimulus, and 2) to which finger it was delivered. Participants answered with their non-stimulated hand using a response box. The next trial began immediately following button press for the second question; there were a total of 50 trials per run. (**B**) Experimental set up. Participants were positioned in a chinrest (to stabilize head position), facing an external monitor (which showed a fixation cross or task-related questions) and an infrared (IR) camera to record eye metrics. The external monitor was attached to a laptop, which ran the tactile task. Signals for the tactile stimuli (sinewaves generated by the laptop) were sent to an amplifier and then to vibrating tactors that were placed on the participant’s fingers. The participant’s free hand was used to control a response box that was connected to the laptop. Signals from the IR camera were sent to a dedicated pupillometry computer. Behavioral task, response, and eye metric data were synchronized via an Ethernet connection. Behavioral task, response, and EEG data were synchronized by TTL pulses, initiated by the laptop, and generated through an Arduino, which was recorded directly through the EEG amplifier.

A computer screen, with a central white fixation cross on a gray background, was placed in front of the participant. The distance from the central fixation cross to the bridge of the participant’s nose was standardized to 55 cm when eye metrics were measured (85 cm when eye metrics were not measured); the size of the displayed screen was adjusted to keep the apparent size and viewing angle (19° across the horizontal dimension) consistent across conditions.

Pre-test training was conducted to familiarize participants with the stimuli. In this training, participants received suprathreshold stimuli to each finger in turn, and were asked to identify which finger had received stimulation. Following training, participants completed four runs of 50 trials each (200 trials total). For each trial (Fig. 1A), participants were asked to fixate on a white cross positioned centrally on a gray background on the computer screen while they waited for a vibration to be delivered to one of their fingers in random order. Participants were told that they may or may not feel a vibration on every trial. Trials began with a randomly jittered 2-4 second period in which the participant fixated on the white cross on the computer screen. Following this period, a 40 ms, 200 Hz vibration was delivered to one of the fingers in 86% of trials; 14% of trials did not have a vibration (blank trials). Stimuli were delivered in random order to the four non-thumb fingers in equal proportions. Following the vibration (or blank), there was an additional 2 or 4 second delay (each occurring 50% of the time) before the first behavioral report question was presented on the screen. Participants were asked two, self-paced forced choice questions, presented successively on the computer screen. The first question (perception question) was: “Did you feel the vibration?” which offered two options: 1 for yes, 2 for no; or 2 for yes, 1 for no. The ‘yes’ button was counterbalanced across participants, but remained constant for the duration of the study for a given participant. Following the perception question, participants were presented with the question (localization question): “On which finger did you feel it?”, with the numbers one to four followed by their corresponding fingers (1-index, 2-middle, 3-ring, 4-pinky). Participants were asked this question regardless of their answer to the first question; if they reported not feeling the vibration, they were instructed to answer the second question randomly. Participants reported their answers to these questions using a response box placed under the hand contralateral to the hand receiving stimulation (Fig. 1B). To aid in answering the second question, both the color and finger identity of the button box corresponded to the hand receiving stimulation (e.g., if they felt the vibration on the ring finger - which had a green foam strap - of the right hand, they should press the ring finger - green button - of their left hand). Data were acquired in runs consisting of 50 trials. Each run took an average of 11.65 minutes, and participants completed a median of 200 trials (range 197 to 250).

### 2.3 Experimental design and equipment

Tactile stimuli consisted of a 200 Hz sinewave pulse (peak sensitivity for Pacinian Corpuscles, (McGlone & Reilly, 2010)) presented for 40 ms. The amplitude of the vibration was titrated in a trial-by-trial manner to approximate the participant’s 50% perceptual threshold, using a minimized expected entropy staircase method (the MinExpEntStair function included in Psychtoolbox, based on Saunders and Backus (2006)). The task was written in MATLAB (The Mathworks Inc., Natick MA, United States) using the Psychophysics Toolbox (‘Psychtoolbox’) extensions (Brainard, 1997; Cornelissen, Peters, & Palmer, 2002; Kleiner et al., 2007; Pelli, 1997). Stimuli were generated in MATLAB, amplified (Marantz NR1609 AV Receiver), and transduced by vibrating tactors (C-2 tactor, Engineering Acoustics, Inc.) placed on the participants’ fingers (Fig 1B).

When eye metrics were measured, participants viewed a fixation cross on a visual display placed directly above a mounted EyeLink 1000 Plus (SR Research, Ottawa, Canada) pupillometer and infrared illuminator. Luminance was controlled across testing sessions by using consistent lighting sources within a windowless testing room. Binocular eye tracking data were collected in head-stabilized mode at 1000 Hz; head stabilization was achieved using a cushioned chin and forehead rest. Prior to the initiation of the behavioral task, participants performed an automated eye-gaze calibration procedure to ensure accurate tracking of eye position.

Non-invasive high-density EEG was recorded from the scalp using a 256-channel net (HydroCel GSN 256, Electrical Geodesics, Inc. Eugene, Oregon). Electrodes were placed using SIGNAGEL (VWR International, LLC, Radnor, PA USA) to enhance conductivity between head and electrodes. After gelling the electrodes, the impedance was measured and was considered acceptable if it was <70 kΩ in more than 90% of the electrodes. Signals were amplified through two 128 channel EEG amplifiers (Electrical Geodesics, Inc. Eugene, Oregon), and recorded and digitized via a NetStation System (1000 Hz sampling rate, high-pass filter of 0.1 Hz, low-pass filter 400 Hz). During recording, channels were Cz-referenced.

Behavioral task, response and pupillometry data were synchronized using digital timing information sent over an Ethernet connection between the behavioral laptop and EyeLink computer, so that the timing of behavioral events (start of trials, runs and questions; stimulus presentations and button presses) could be recorded on the same time base as the EyeLink recordings. In addition, to ensure precise synchronization between behavioral task, response, and EEG data, TTL pulses from the behavioral laptop were directly input to the EEG amplifier and recorded as event flags on the EEG recording on Netstation. TTL pulses – again corresponding to the start of trials, runs, and questions; stimulus presentations and button presses – were initiated by the laptop (Macbook Pro) running the task and generated by an Arduino Uno (R3; Smart Projects) that connected to the digital input port of the EEG amplifier via a DB9 cable. Responses were recorded using a four-button response box (Current Designs, Inc., Model OTR-1×4-L), which was connected to the laptop via USB and sampled by the computer at 1000 Hz.

### 2.4 Data Analysis

#### 2.4.1 Behavioral analysis

Trials were considered for analysis if they were classified as confirmed perceived, or confirmed not perceived, validated by the location question. Trials in which a vibration was present, reported as felt, and then localized to the correct finger were considered confirmed (validated) perceived; trials in which a vibration was present, reported as not felt, and then incorrectly localized were considered confirmed (validated) not perceived (Fig. 2B).

**Figure 2:**
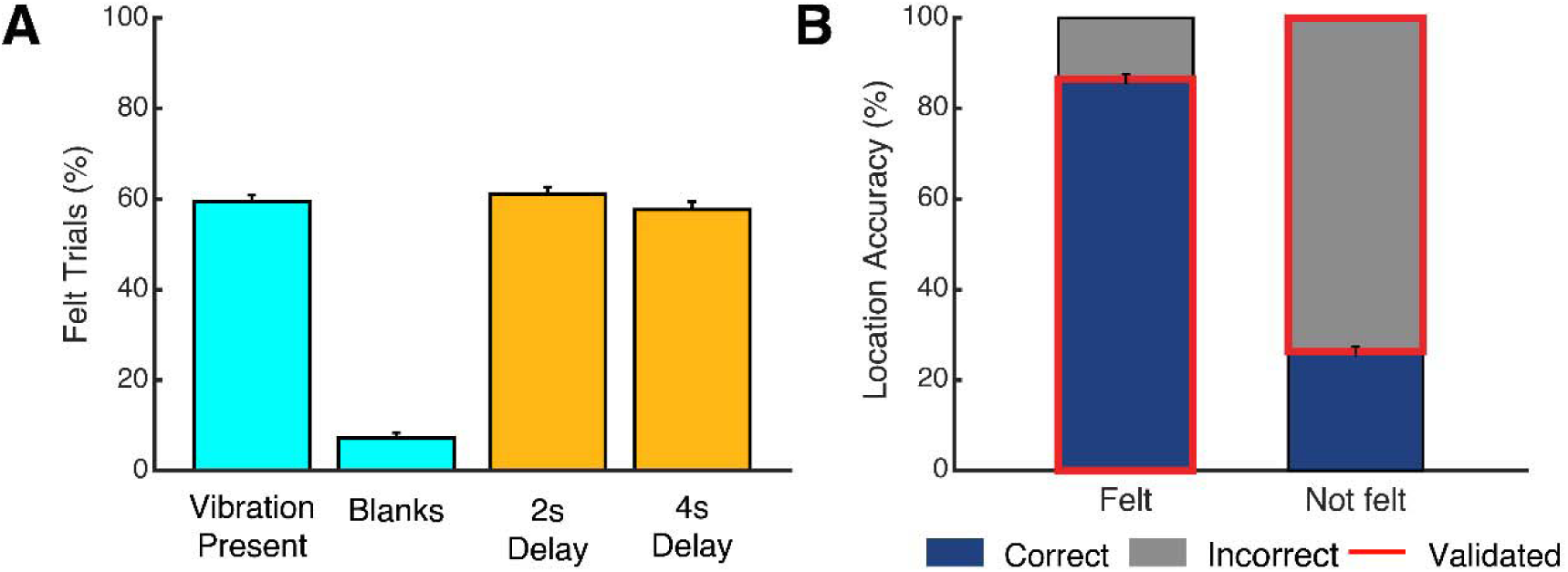
Behavioral results. **(A)** Responses to perception question. In trials in which a vibration was present, 59% were reported as felt; in trials in which no vibration was played, only 8% were reported as felt. Report of vibration was approximately the same for 2 or 4 s post-stimulus delays. Error bars are standard error of the mean (SEM). **(B)** Responses to location question. When vibrations were reported as felt, participants correctly reported which finger received a vibration for 86% of trials; when they reported they did not feel a vibration (in trials when there was a vibration present), they reported the finger incorrectly for 73% of trials (chance=75%). Correctly identified trials are shown in navy blue; incorrectly identified trials are shown in gray. Data considered for analysis are highlighted in red. Error bars are SEM.

Because the tactile perceptual threshold was observed to change across the course of a single behavioral session, a continually adjusting staircase method was used to approximate the instantaneous perceptual threshold. This method results in many trials that are at threshold, but some that are presented at amplitudes supra- and sub-threshold; therefore, a Euclidean distance analysis was used to select trials that were presented closest to the perceptual threshold, and to match the magnitude of confirmed perceived and confirmed not perceived trials used in the analysis. To do so, for each participant, on a finger-by-finger basis, trials categorized as confirmed perceived and confirmed not perceived were selected. The order of confirmed perceived trials was randomized; the shuffled order of confirmed perceived trials was then used to match each confirmed perceived trial’s amplitude to the confirmed not perceived trial with the closest amplitude. If the amplitude difference between the confirmed perceived and confirmed not perceived trials fell within 0.03 (arbitrary units [au]; the testing algorithm could adjust in 0.001 au increments), the pairing was included and both trials were removed from their respective pools; if it fell outside of those boundaries, that confirmed perceived trial was discarded and the confirmed not perceived trial was replaced into the not perceived pool. This continued until all confirmed perceived trials were either paired with a unique confirmed not perceived trial or discarded. The total number of trials included was tallied, and the sum of differences between each unique pairing was calculated. After 100,000 replications of this procedure, the replication with the largest number of trials was retained, and the trails from that replication were selected for analysis. If two or more replications yielded the same number of included trials, the replication with the smallest sum of amplitude differences was selected; if this was also identical, a replication was chosen randomly from the equivalent replications. For simplicity, we will hereafter refer to this selected subset of trials as perceived and not perceived, with the understanding that all analyzed trials have been validated by both localization accuracy (see the preceding paragraph) and proximity to the participant’s perceptual threshold (current paragraph).

#### 2.4.2 Event-Related Potential (ERP) analysis

After extraction from the NetStation system, the EEG data were analyzed using MATLAB and EEGLAB (Delorme & Makeig, 2004). For each participant, a high-pass 0.1 Hz filter and the CleanLine procedure (Mullen, 2012) were applied to exclude low-frequency drifts and line noise in the 60 Hz and 120 Hz frequency bands. To reject trials with high-frequency noise (e.g., from muscle or movement artifact), high frequency power was calculated using a high-pass 30 Hz filter applied at the channel level across the entirety of the session (window size: 4 s, 2 s overlap). Epochs were identified and cut from -2,000 ms to +2,000 ms, centered on the vibration onset. All further analyses were conducted independently for trials belonging to either perceived or not perceived trials (see 2.4.1). Channels with excessive high frequency power (20% or more timepoints of the filtered high frequency power in a trial exceeded 100μV) were excluded, and their positions were re-populated using a spherical interpolation (EEGLAB pop_interp function, there were no more than 10% of channels deleted in a trial). The resulting data were re-referenced to the average of the mastoids’ signals. Epochs were collated and passed through a semi-automatized principal component analysis (PCA) and Independent Component Analysis (ICA) decomposition rejection procedure (EEGLAB pop_runica function utilizing the infomax algorithm for ICA decomposition), in which the ten principle components that explained the most variance of the data were identified, and then among these components 10 independent components were found. Trained study personnel removed independent components that corresponded to signatures for blink, eye-movement, and heartbeat artifacts. Finally, a 25 Hz lowpass filter was applied, and the average of perceived, and not perceived epochs were acquired. The resulting signals were baselined by subtracting the mean of the interval from -1000 to -1 ms (the second preceding stimulus).

To control for effects of lateralization, the brain maps of participants that received the vibrations on their left hand were mirrored, with the electrodes assuming the position of their contralateral equivalents. Therefore, for all analyses, the electrodes on the left side of the head represent signals contralateral to the side of the stimulated hand, and electrodes on the right side of the head represent signals ipsilateral to the side of the stimulated hand. After pooling data by first calculating within-participant mean ERPs, the means and SEM were calculated across participants. The results were then resampled at 200 Hz (using MATLAB function imresize) and re-baselined by subtracting the pre-stimulus period of -1000 to -5 ms. False discovery rate (FDR) analyses were applied to 0 to 1000 ms post-stimulus to identify areas of significance (null hypothesis: voltage=0 µV, q<0.05) in the grand average ERPs. This procedure aimed to control for multiple comparisons using the mass univariate analysis (MUA) and EEGLAB’s ERPLab toolboxes (Benjamini & Hochberg, 1995; Benjamini & Yekutieli, 2001; Groppe, Urbach, & Kutas, 2011; Lopez-Calderon & Luck, 2014). This FDR procedure controls for multiple comparisons across electrodes and timepoints by computing the p-values for each timepoint at each electrode, combining, and sorting the entire distribution of p-values, and computing a threshold based on alpha level. For the P300 peak times, the timepoint with the highest voltage for the Pz electrode was found for the average ERP across participants.

#### 2.4.3 Eye metrics analyses

Eye-metric data were analyzed in custom software written in MATLAB. First, to prepare eye-metric data for analysis, artifact rejection was conducted to remove invalid portions of data. Blinks and artifacts were detected by implementing a MATLAB procedure called Stublinks (Kronemer et al., 2021; Greg J. Siegle, Steinhauer, Stenger, Konecky, & Carter, 2003). Data segments were flagged if no pupil was detected (due to blink or loss of signal); or if signal spikes were detected (e.g., those associated with the opening or closing of the eyelid during a blink, or those differing more than 4 mm from a trial’s median diameter). Segments of flagged data that lasted from 100-400 ms were labeled as blinks based on their duration (Schiffman, 2001) and used to generate the blink timecourse data; other flagged segments were marked as artifact. For pupil diameter and gaze (microsaccade) analyses, the rejected samples were linearly interpolated (MATLAB stublink function for pupil data, and naninterp for gaze data) with temporally adjacent samples to restore the omitted time points.

Eye metrics (pupil diameter, blink rate, microsaccade rate) were analyzed as a function of trial type (e.g., perceived or not perceived) on a per participant basis, and then averaged across participants. For each metric, a time window from 1000 ms before the vibration to 2000 ms following vibration onset was extracted and analyzed.

To calculate the mean pupil diameter timecourse, we first baseline-corrected the data to control for changes in steady-state (e.g., not event-related) pupil diameter across runs, or differences across participants. This was achieved by subtracting the median pupil diameter from the 1000 ms preceding the onset of the vibration on a trial-by-trial basis. The mean of the resulting baseline-corrected timecourses was calculated within trial condition (e.g., perceived or not perceived) within each participant; the grand mean across participants was then calculated.

Blink rate, using the detected blinks, corresponds to the proportion of trials that had a blink occurring at a given time point (e.g., if 20 out of 100 trials had a blink occurring during time *t*, the blink rate at time *t* would be 0.2). Blink rate was calculated for each sample; no binning or baselining was applied. The blink rate was done on a participant level, and then averaged across participants.

Saccades were extracted from the eye tracking data and the ones smaller than one degree (microsaccades) were identified using the algorithm described by Engbert and Kliegl (2003). Microsaccade rate was calculated by identifying the number of saccades initiated inside 500 ms windows (successive windows overlapped by 250 ms). On a trial basis, the number of microsaccades initiated within a given window were tallied; this was then converted to the rate of microsaccades per second (e.g., if 3 microsaccades were initiated, the rate within that 500 ms window would be 6 microsaccades/second or 6 Hz). Mean microsaccade rates were calculated across trials for each participant; these means were then used to calculate a grand mean of microsaccade rate across participants.

To calculate when pupil diameter, blink rate, and microsaccade rate significantly differ as a function of perception, we performed a bootstrap analysis. First, the grand mean of the not perceived trials was subtracted from the grand mean of the perceived trials for each eye metric. For each bootstrap, trials were randomly selected (with replacement) from the original dataset, and given a randomly shuffled perceived or not perceived trial label. These relabeled trials were analyzed in the same manner as the original data. The resulting group-averaged timecourses from bootstrapped trials assigned to the “not perceived” group was subtracted from that of bootstrapped trials assigned to the “perceived” group. This procedure was repeated 10,000 times. The grand mean of the 10,000 bootstrapped, subtracted timecourses, and confidence intervals at each time point were then calculated. Timepoints (or bins) in the original data were considered to have a significant difference between perceived and not perceived conditions if there was a less than 5% chance of the observation occurring in the bootstrapped data.

## 3 Results

### 3.1 Behavioral results

Participants reported feeling a vibration in 59 ± 1% (mean ± SEM) of trials in which there was a vibration present; this remained relatively consistent across the two post-stimulus delays (2 seconds: 61 ± 1%; 4 seconds: 57 ± 2%); only two participants had significant differences in their percentage perceived as a function of post-stimulus delay (chi-square<0.05)) (Fig 2A). Participants reported feeling a vibration on only 8 ± 1% of the blank trials (Fig 2A). On average, 86 ± 1% of the trials reported as felt were also reported in the correct location (Fig 2B); 27 ± 1% of trials that were reported as not felt were reported on the correct finger (chance is 25%). After the Euclidean distance analysis (see Methods), an average of 44 ± 8 trials per condition were included per participant for analysis.

### 3.2 Evoked potentials

At early times, a prominent N140 was observed bilaterally in frontal areas for perceived trials (Fig. 3). It reached significance (q < 0.05, see Methods) for both perceived analysis alone and when comparing the difference between perceived and not perceived trials. For the not perceived condition, a significant (q<0.05) P100 was found bilaterally in parietal and occipital areas (Fig. 3 and S1). Significant findings remain the same when the data were not mirrored (i.e., when stimulated hand was not controlled for; see S2).

**Figure 3:**
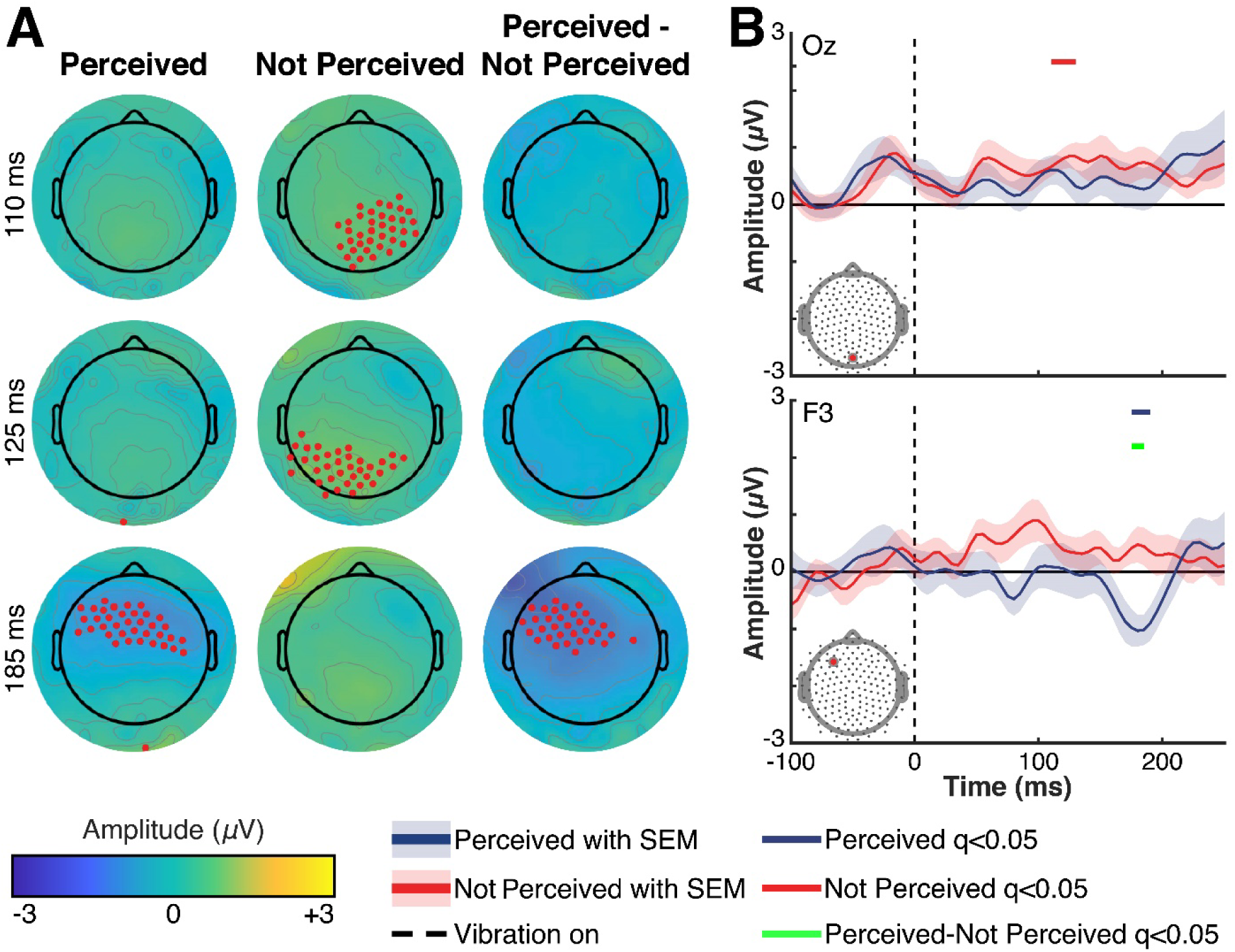
Early ERPs. (**A**) Voltage topographic maps for early ERPs. Electrodes that achieve significance using an FDR analysis (data analyzed: 0 to 1000 ms post vibration, null hypothesis: voltage=0 µV, q<0.05) are highlighted in red. Times are relative to vibration onset, and were chosen to highlight specific signals of interest (see Fig. S1 for maps at all times). (**B**) Voltage timecourses highlighting early changes post-vibration (−100 ms to +250 ms from vibration onset). Electrode positions are indicated via a red marker on the map on the bottom right corner of each plot. Blue traces show timecourses of perceived trials; red traces show timecourses of not perceived trials. Shaded error bars show respective SEMs. Red, blue, and green lines at the top of each plot indicate windows that reached significance using FDR methods. Blue corresponds to significant windows in perceived data; red for not perceived data; and green for the perceived-not perceived data. Times are relative to the onset of vibration, represented by the vertical dotted line.

At later times, significant P300 responses were found for both perceived and not perceived trials; however, the spatial extent, duration, and magnitude were larger for perceived trials (Fig. 4, S1). For perceived trials, the P300 reached peak at 290 ms post-vibration onset, but showed significance above baseline from 260-855 ms post-vibration. In contrast, the P300 for not perceived trials has an overall peak at 440 ms (Fig. 4, S1).

**Figure 4:**
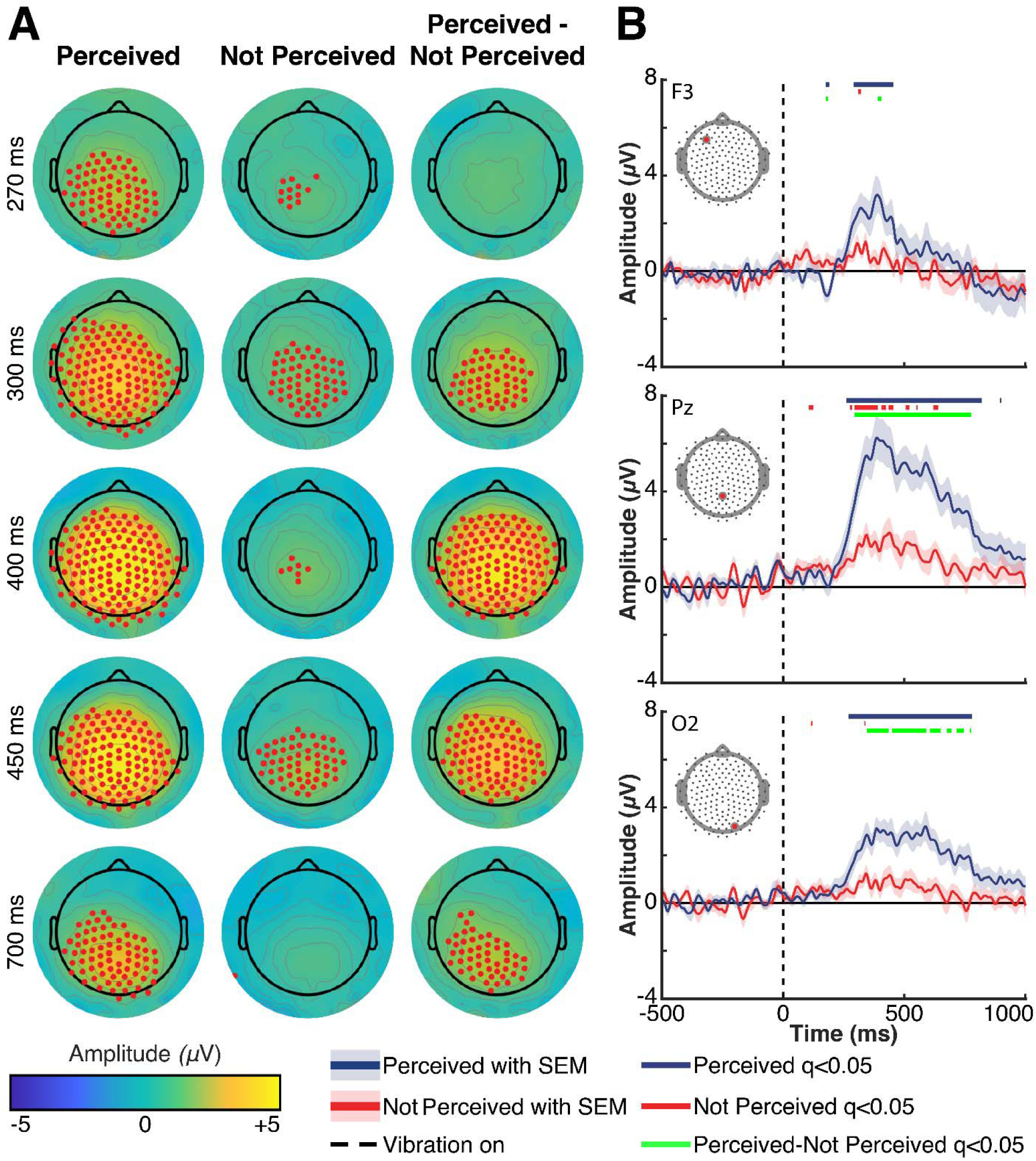
Late ERPs. (**A**) Voltage topographic maps for late ERPs. Electrodes that achieve significance using an FDR analysis (data analyzed: 0 to 1000 ms post-vibration, null hypothesis: voltage=0 µV, q<0.05) are highlighted in red. Times are relative to vibration onset, and were chosen to highlight specific signals of interest (see Fig. S1 for maps at all times). (**B**) Voltage timecourses highlighting late changes post-vibration (−500 ms - +1000 ms post-vibration onset). Electrode positions are indicated via a red marker on the map on the bottom right corner of each plot. Blue traces show timecourses of perceived trials; red traces show timecourses of not perceived trials. Shaded error bars show respective SEMs. Red, blue, and green lines at the top of each plot indicate windows that reached significance using FDR methods. Blue corresponds to significant windows in perceived data; red for not perceived data; and green for the perceived-not perceived data. Times are relative to the onset of vibration, represented by the vertical dotted line.

### 3.3 Eye metrics

Significant differences were found between perceived and not perceived conditions for pupil diameter, blink rate, and microsaccade rate. Pupil diameter increased markedly for perceived trials, peaking on average approximately 1100 ms following a vibration (Fig. 5A). Pupil diameter was significantly different between perceived and not perceived trials from ∼270 ms after vibration to the end of the analyzed epoch (2000 ms post-vibration onset). Blink rate also increased following a vibration for perceived trials, reaching a peak rate ∼800 ms post-vibration. Blink rate differed significantly between perceived and not perceived trials for most of the period from ∼500 ms post-vibration to the end of the analyzed epoch (Fig. 5B). While pupil diameter and blink rate showed an increase for perceived trials relative to not perceived trials, the microsaccade rate dynamic was more complicated: microsaccade rate was suppressed in perceived trials relative to not perceived trials from ∼250-1000 ms post-vibration; but then significantly increased above not perceived microsaccade rate from 1250 ms post-vibration until the end of the analyzed epoch (Fig. 5C).

**Figure 5:**
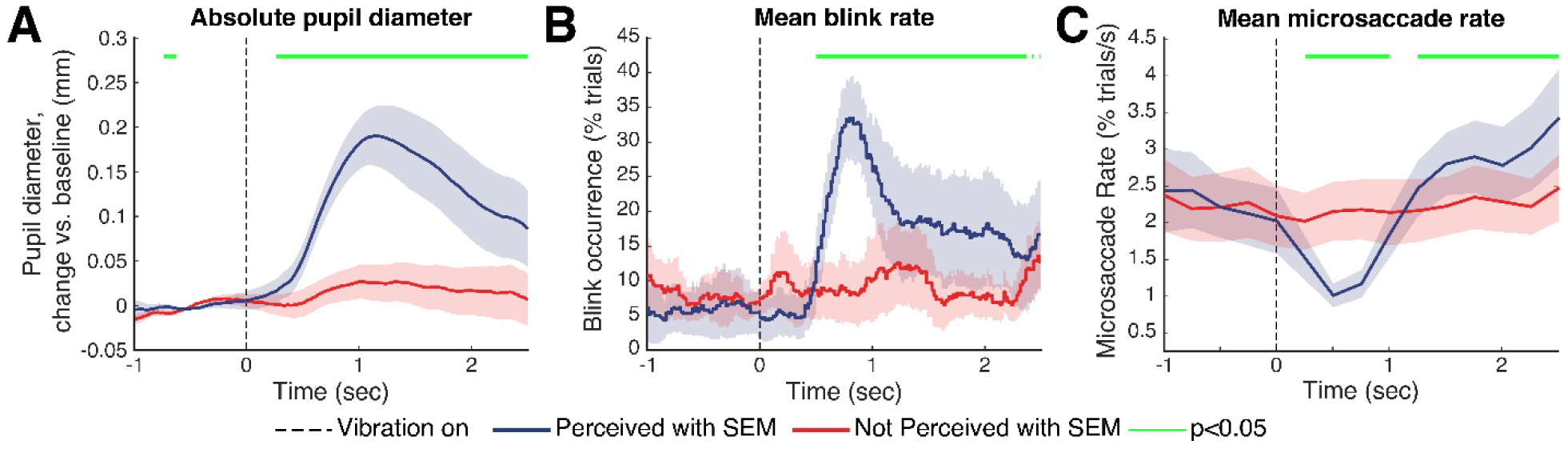
Eye metrics. Timecourses of (**A**) average pupil diameter (**B**) mean percentage of trials where there is a blink occurrence; and (**C**) mean microsaccade rate per second. Blue traces represent the grand mean of perceived trials; red traces represent the grand mean of not perceived trials. Shaded error bars show respective SEMs. Times for which there is a significant difference between perceived and not perceived trials are indicated by the green line at the top of each plot. Times are relative to the onset of vibration, represented by the vertical dotted line.

## 4 Discussion

In this study, we investigated the ERP and eye metrics correlates of tactile conscious perception using mechano-vibrational stimuli in a threshold task. For perceived stimuli, we found significant ERPs corresponding to the N140 and P300; for not perceived trials, we observed significant ERPs for the P100 and P300. We also found a significant N140 and P300 when comparing the difference in signals between perceived and not perceived trials (Fig. 3 and 4, S1). Finally, we noted an increase in pupil diameter and blink rate, and a decrease in microsaccades following perceived vibrations (Fig. 5).

These findings are not without precedent. Perhaps the best documented perception-related ERPs are the components of the P300, found in this study starting ∼260 ms and lasting until 855 ms. The P300 has been variously attributed to allocation of attentional resources (Donchin & Coles, 1988; Muñoz et al., 2014), awareness and conscious perception, or to perceptual report or post-perceptual processing (Cohen et al., 2020; Dehaene & Changeux, 2011; Del Cul et al., 2007; Kronemer et al., 2021; Pitts, Padwal, Fennelly, Martinez, & Hillyard, 2014). Although it has historically been considered a marker of consciousness, multiple papers using different paradigms have more recently shown that the P300 is not present in the absence of perceptual report or when a suprathreshold stimulus is not task-relevant (Cohen et al., 2020; Derda et al., 2019; Kronemer et al., 2021; Pitts, Metzler, et al., 2014; Pitts, Padwal, et al., 2014; Polich, 2007; Railo et al., 2011; Ye et al., 2019). Our task does require perceptual report — even in the absence of a vibration or perception, and so it is interesting to note that we observe a P300 in both perceived and not perceived trials. However, the amplitude and spatial extent of the P300 is greater for perceived trials. Pitts, Metzler, et al. (2014) also found a smaller positivity at ∼300ms for not perceived that they called P3b, but – as in our data – it had a smaller amplitude than the one found for perceived trials. Because of that, they believe that the P300 is likely to reflect post-perceptual or attention-based processes necessary for completing the task.

The other signals we observed also have precedence in perception literature. Previous studies have shown the P100 and N140 in relation to perceived or salient stimuli; these signals were stronger in the absence of masking or surrounding stimuli (Kida et al., 2006; Schubert et al., 2006). Schubert et al. (2006) suggest that the P100 marks the emergence of perception for a perceived stimulus. Because we find a P100 for not perceived trials but not for perceived, our data is inconsistent with the attribution of the P100 as simply a marker of positive perceptual status.

Dembski, Koch, and Pitts (2021) suggest that the N140 negativity is the somatosensory equivalent to the visual and the auditory awareness negativity (VAN and AAN respectively), and that these negativities, collectively, should be considered a main marker of consciousness. However, there is some variability about the interpretation and localization of a negativity occurring at ∼140 ms post-stimulus across studies. Dembski et al. (2021) hold that because the VAN and AAN are localized to their respective primary processing areas, the somatosensory awareness negativity (SAN) should be similarly located. This is supported by the findings of Auksztulewicz and Blankenburg (2013) and Forster et al. (2016) who each find a central-contralateral negativity between 100-200 ms. Notably, Forster et al. (2016) interpret their N140 as a measure of spatial attention, not awareness, but their study was not designed to test conscious awareness. In contrast, Schubert et al. (2006) report the N140 bilaterally in frontal (not central) electrodes, but also suggest that it is linked to conscious awareness and is modulated by spatial attention. Our N140 is observed bilaterally in frontal-central electrodes — like Schubert et al. (2006) — but with somewhat greater involvement of contralateral locations. As the N140 is significant for perceived trials, and when comparing perceived to not perceived trials, we suggest that it should, indeed, be considered a marker of conscious awareness. The greater involvement of contralateral electrodes may be due to the allocation of spatial attention to the stimulated hand, and therefore is also in potential agreement with Auksztulewicz and Blankenburg (2013) and Forster et al. (2016).

A fundamental challenge of conscious perception research is distinguishing brain activity associated with pre- or post-perceptual processing—including perceptual report. The need to disambiguate perceptual report, specifically, from perception is the center of field-wide divisions on theories of consciousness. Indeed, the interpretation of what the P300 represents hinges partially on whether it is a marker of consciousness or a result of report. To remove the potential confounds of using either differential stimuli (such as masks), recent work has explored eye metrics as a covert measure of perception that may open the door for the development of no-report paradigms (Babiloni et al., 2001; Cohen et al., 2020; Derda et al., 2019; Donchin & Coles, 1988; Egsgaard et al., 2012; Koivisto, Grassini, Salminen-Vaparanta, & Revonsuo, 2017; Koivisto et al., 2016; Muñoz et al., 2014; Pitts, Metzler, et al., 2014; Pitts, Padwal, et al., 2014; Polich, 2007; Railo et al., 2011; Truini et al., 2004).

Although eye metrics – especially pupil diameter – are widely used in visual perception studies (Aminihajibashi, Hagen, Laeng, & Espeseth, 2020; Aston-Jones & Cohen, 2005; Choe, Blake, & Lee, 2016; Eckstein et al., 2017; Geng, Blumenfeld, Tyson, & Minzenberg, 2015; Otero-Millan, Macknik, Serra, Leigh, & Martinez-Conde, 2011; Wang, Blohm, Huang, Boehnke, & Munoz, 2017) and in some auditory ones (Wetzel, Buttelmann, Schieler, & Widmann, 2016; Zekveld, Koelewijn, & Kramer, 2018), studies using tactile stimuli are still scarce (Gusso, Serur, & Nohama, 2021). To our knowledge, there are no prior reports of eye metrics in threshold tactile perception tasks; and in most other tactile studies, stimuli were delivered manually (Iriki, Tanaka, & Iwamura, 1996; van Hooijdonk et al., 2019) or used recording systems with a sampling rate that, according to Holmqvist et al. (2011), is insufficient to capture nuanced physiological changes occurring at the eye-level (Bertheaux et al., 2020; Ganea et al., 2020; Iriki et al., 1996; C. C. Y. Lee, Kheradpezhouh, Diamond, & Arabzadeh, 2020; C. R. Lee & Margolis, 2016; Schriver, Bagdasarov, & Wang, 2018; Schriver, Perkins, Sajda, & Wang, 2020; van Hooijdonk et al., 2019). The ideal sampling rate for eye metrics, especially if microsaccades are to be measured, is >> 250 Hz; no mathematical method defines a cut-off value, but it instead has been established through consensus in order to be able to acquire all physiological changes occurring on the eye-level (Holmqvist et al., 2011).

We could not find studies associating blink rates to tactile stimuli. However, previous studies have shown that blink rate is inversely related to cognitive or attentional demand: blink rate decreases with increased attention, and increases when attentional or cognitive demands are removed (Fukuda & Matsunaga, 1983; Greg J Siegle, Ichikawa, & Steinhauer, 2008). The increase in blink rate immediately following a perceived vibration is consistent with these findings; once a vibration has been felt, the participant has ‘achieved’ their goal and no longer needs to closely attend to the trial.

Microsaccades have been associated with visual accommodation and a necessary physiologic response so we can process the visual world (Otero-Millan et al., 2011). Here, we show that in addition to being affected by visual paradigms, changes in microsaccade rate can be elicited by tactile perception—notably, by a decrease in microsaccades after perception. This decrease in rate is consistent with a recent study from Badde et al. (2020), who reported oculomotor freezing after cue acquisition. Although our study does not use cues, both Badde et al. (2020) and our current findings are consistent with decreased involuntary eye movement after a perceived sensory event, independent of that event’s modality.

The pupil, blink and microsaccade changes we observed with the tactile threshold perception task were very similar to those recently reported for a similarly-designed visual threshold perception task (Kronemer et al., 2021). The consistency of eye metrics in perceptual tasks across sensory modalities, now also encompassing tactile perception, lends further support to the idea that eye metrics serve as a robust covert measure of electrophysiological changes associated with cognitive engagement and its associated changes in physiological arousal levels. The similarity of eye-metric dynamics across sensory modalities and paradigms suggests that eye metrics represent a potentially powerful tool for gauging perceptual and cognitive processing in the absence of overt perceptual report. This approach has recently been applied successfully to conscious visual perception (Kronemer et al., 2021). We plan to leverage these metrics in the development of no-report paradigms in future studies across sensory modalities.

## 5 Conclusions

Overall, our current study uses a novel tactile threshold paradigm combined with high-density scalp EEG, pupillometry and eye-tracking. We report, for the first time using a tactile-threshold task, that ERPs similar to those often associated with perceived stimuli in other sensory domains, such as the N140 and P300, are elicited by perceived tactile stimuli. We note that the P300 (of lower amplitude) is also elicited in our not perceived trials, further complicating the already complex story of what the P300 may represent. We also present, for the first time, that pupil diameter, microsaccade rate, and blink rate differ in a tactile threshold perception task, which suggests that eye metrics may represent a path toward the creation of tactile no-report tasks in the future.

## Abbreviations

FDR: false discovery rate
MUA: mass univariate analysis

## Authors’ Statement

**Mariana M. Gusso**: Conceptualization, Methodology, Software, Formal analysis, Investigation, Writing – Original Draft, Visualization. **Kate L. Christison-Lagay**: Conceptualization, Methodology, Software, Formal analysis, Writing – Original Draft, Writing – Review & Editing, Visualization. **David Zuckerman**: Investigation. **Ganesh Chandrasekaran**: Investigation. **Sharif I. Kronemer**: Methodology, Formal Analysis, Writing – Review and Editing. **Julia Z. Ding**: Formal Analysis. **Noah C. Freedman**: Formal Analysis. **Percy Nohama**: Methodology. **Hal Blumenfeld**: Conceptualization, Methodology, Resources, Data Curation, Writing – Review and Editing, Supervision, Project administration, Funding Acquisition.

## Acknowledgements

We thank Yale’s Child Study Center for use of research space, and Dr. Michael Pitts (Reed College) for his advice on analytical approaches.

## Supplemental labels

**Supplemental material S1: Voltage topographic maps timecourse (movie)**. Voltage topographic maps from 0-1000 ms. Electrodes that achieve significance using an FDR analysis (null hypothesis: voltage=0 microvolts, q<0.05) are highlighted in red. Times are relative to vibration onset.

**Supplemental material S2:**
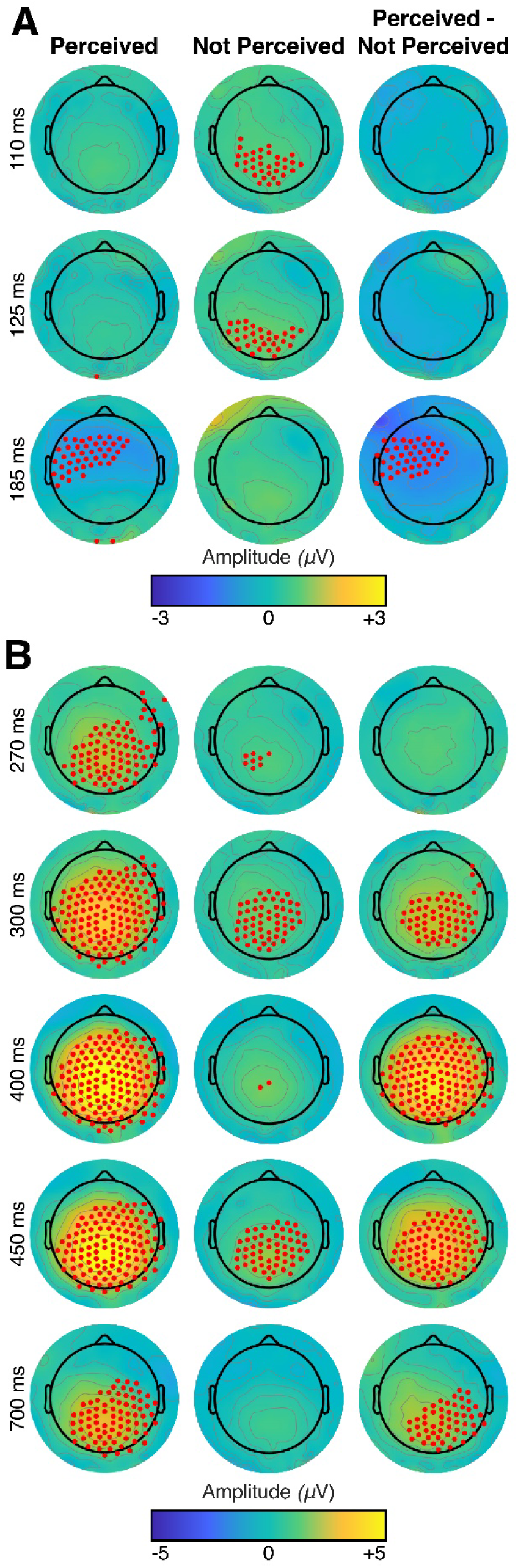
Voltage topographic maps without mirroring. Maps are shown without mirroring the maps for the participants who received the stimuli on the left hand (see Methods) for (**A**) early ERPs (0-250 ms post vibration), and (**B**) late ERPs (250-700 ms post vibration). Electrodes that achieve significance using an FDR analysis (null hypothesis: voltage=0 microvolts, q<0.05) are highlighted in red. Times are relative to vibration onset, and were chosen to highlight specific signals of interest.

